# Specialized mechanoreceptor systems in rodent glabrous skin

**DOI:** 10.1101/337923

**Authors:** Jan Walcher, Julia Ojeda-Alonso, Julia Haseleu, Maria K. Oosthuizen, Ashlee H. Rowe, Nigel C. Bennett, Gary R. Lewin

## Abstract

Rodents use their forepaws to actively interact with their tactile environment. Studies on the physiology and anatomy of glabrous skin that makes up the majority of the forepaw are almost non-existent in the mouse. Here we developed a preparation to record from single sensory fibers of the forepaw and compared anatomical and physiological receptor properties to those of the hind paw glabrous and hairy skin. We found that the mouse forepaw skin is equipped with a very high density of mechanoreceptors; >3 fold more than hind paw glabrous skin. In addition, rapidly adapting mechanoreceptors that innervate Meissner’s corpuscles of the forepaw were several-fold more sensitive to slowly moving mechanical stimuli compared to their counterparts in the hind paw glabrous skin. All other mechanoreceptors types as well as myelinated nociceptors had physiological properties that were invariant regardless of which skin area they occupied. We discovered a novel D-hair receptor innervating a small group of hairs in the middle of the hind paw glabrous skin in mice. Glabrous D-hair receptors were direction sensitive albeit with an orientation sensitivity opposite to that described for hairy skin D-hair receptors. Glabrous D-hair receptors do not occur in all rodents, but are present in North American and African rodent species that diverged more than 65 million years ago. The function of these specialized hairs is unknown, but they are nevertheless evolutionarily very ancient. Our study reveals novel physiological specializations of mechanoreceptors in the glabrous skin that likely evolved to facilitate tactile exploration.

## Introduction

In the past the tactile sense of rodents has been investigated predominantly through the study of hairy skin sensation (Lechner and Lewin, 2013; Li et al., 2011). This is despite the fact that many classical behavioral assessments of rodent sensation such as the Hargreaves test (Hargreaves et al., 1988), or mechanical withdrawal threshold measurements with von Frey hairs are actually carried out by stimulating the glabrous skin (Peng et al., 2017; Ventéo et al., 2016; Wetzel et al., 2017). In addition, rodents constantly use their forepaws to explore their environment, for example selecting food objects or engaging in grooming behavior. Indeed such exploratory or active touch tasks uniquely involve the forepaw glabrous skin as the primary sensory surface used. The mouse forepaw is in many respects analogous to the human hand, but has hardly been examined at the functional or anatomical level. A more detailed exploration of glabrous skin sensory receptors in the mouse is even more relevant considering the increasing interest in skilled forelimb movements, tactile feedback during movement and perceptual tasks based on sensory stimuli applied to the forepaw glabrous skin (Estebanez et al., 2017; Fink et al., 2014; Milenkovic et al., 2014; Wetzel et al., 2017). Several types of mechanoreceptors have been characterized using electrophysiology mostly in the mouse hairy skin(Koltzenburg et al., 1997; Lewin and Moshourab, 2004; Milenkovic et al., 2008). One mechanoreceptor type important for tactile sensation is the rapidly adapting mechanoreceptor (RAM) which fires only to skin movement, but innervates morphological distinct end-organs in hairy skin and glabrous skin (Lewin and Moshourab, 2004; Omerbašić et al., 2015). Thus RAMs innervating hairy skin form lanceolate endings around hair follicles, but innervate Meissner’s corpuscles in the glabrous skin (Heidenreich et al., 2012; Li et al., 2011; Wende et al., 2012). However, even at the molecular level RAMs in glabrous and hairy skin utilize the same potassium channel KCNQ4 to regulate their sensitivity to low frequency vibratory stimuli (Heidenreich et al., 2012). The functional properties of hairy and glabrous skin RAMs are thought to be similar in humans, but this has not been systematically investigated in rodents.

The primary aim of the present study was to investigate the physiology and anatomy of cutaneous afferents in the mouse forepaw skin. In order to answer the question of whether forepaw afferents differ significantly from those in other skin regions we also used identical methods to record from hind paw glabrous and hairy skin receptors. Here we used the *ex vivo* skin nerve method, a well-established technique for studying single sensory receptors in rodents. The most commonly used preparation is that of the saphenous nerve, which innervates the hairy skin of the lateral foot and ankle (Koltzenburg et al., 1997; Kress et al., 1992; Reeh, 1986). Only a few studies have recorded from sensory receptors in the tibial nerve in rodents, which innervates the hind paw glabrous skin (Cain et al., 2001; Leem et al., 1993; Milenkovic et al., 2014). Here we developed a novel *ex vivo* skin nerve preparation to record from mouse sensory receptors with axons in the ulnar and median nerves which innervate the forepaw glabrous skin. We compared data using this preparation with recordings from sensory receptors in the tibial and saphenous nerves that innervate hindlimb glabrous and hairy skin, respectively. We show that RAMs that innervate Meissner’s corpuscles in the forepaw skin are much more sensitive to low frequency vibration stimuli and are also present at a much higher density than those in other skin areas. Myelinated nociceptors had physiological properties that were invariant across skin areas. We also discovered a novel D-hair receptor population innervating a distinct group of very small hairs only found within the hindlimb glabrous skin. These glabrous D-hair receptors are not ubiquitous in rodents, but do occur in rodent species from North America and Africa that diverged more than 65 million years ago (Fabre et al., 2012). Thus our study reveals unique features of both forepaw and hind paw glabrous skin receptors that are highly relevant for tactile driven behavior.

## Results

### Comparative study of sensory afferents innervating forepaw and hind paw

We directly compared the functional properties of identified mechanoreceptors and myelinated nociceptors across three skin areas of the mouse. We used an established *ex vivo* preparation for the hind paw hairy and glabrous skin innervated by the saphenous and tibial nerves, respectively (Koltzenburg et al., 1997; Milenkovic et al., 2008, 2014) (Fig. 1A-F). In addition, we established a new *ex vivo* preparation that enabled us to record from the median and ulnar nerves that predominantly innervate the mouse forepaw glabrous skin (see Methods and Fig. 1G-I). We used teased nerve fiber recordings to record from single-units with myelinated axons and characterized their receptor properties in detail. We took a comparative approach using identical methodologies to record from and characterize the receptor properties of all myelinated afferents forming endings in the hind paw and forepaw skin. In total, we made recordings from 111 Aβ-fibers (conduction velocities >10m/s) and 92 Aδ-fibers (conduction velocities between 1.0 and 10m/s) (data summarized in Table 1). Single-unit data was obtained from 52 mice.

**Figure 1:**
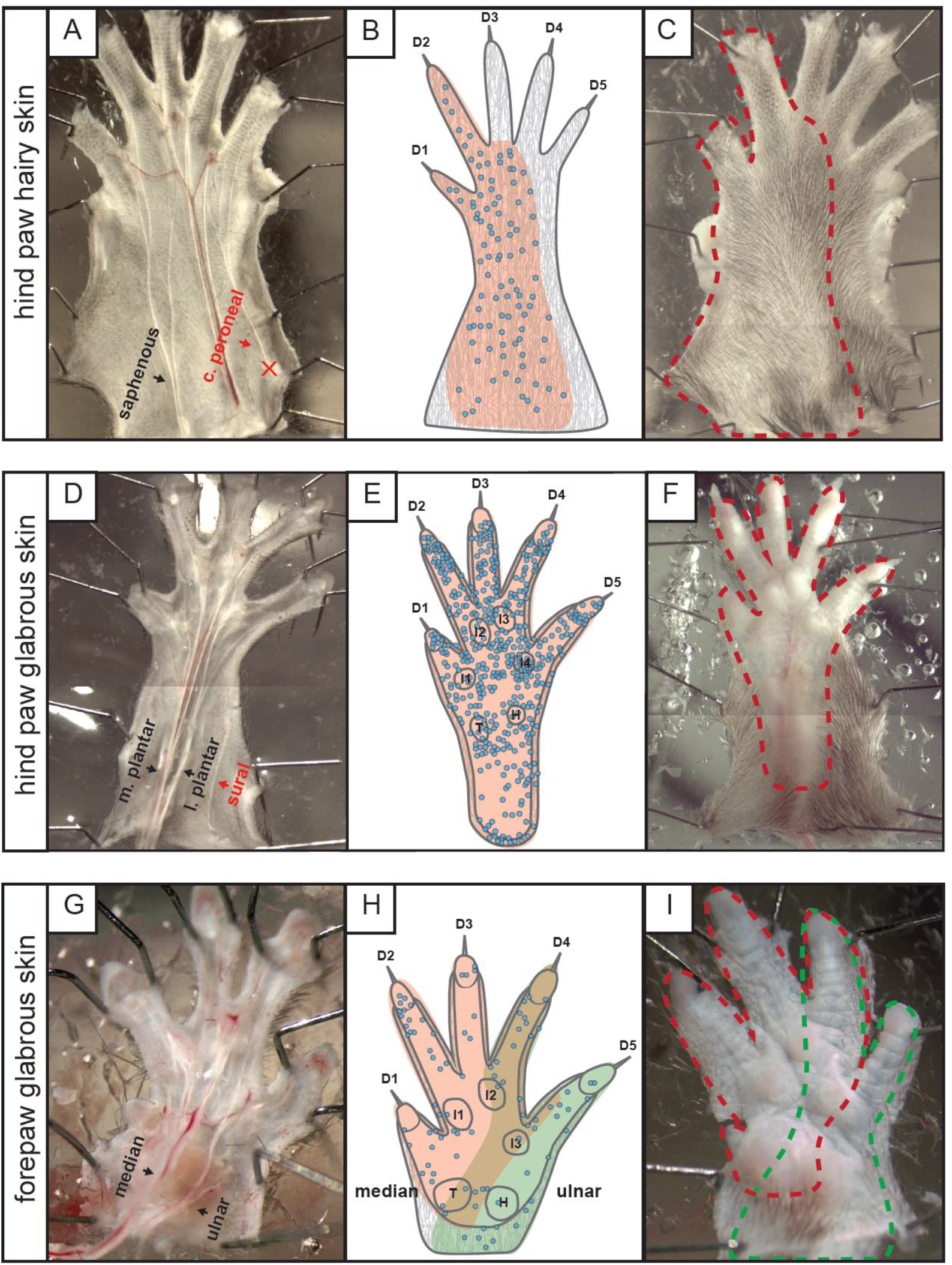
Innervation areas of the saphenous, tibial and median/ulnar nerves. *Left:* Inside out configuration of **A** the hind paw hairy skin, **D** the hindpaw glabrous skin and **G** the forepaw glabrous skin preparation. Nerves indicated in red were not used and the location of the cut end is marked with (X) *Middle:* Receptor locations of single units recorded from **B** the saphenous nerve, **E** the medial and lateral plantar nerve which are two branches of the tibial nerve and **H** the median (red area) and ulnar nerves (green area) are indicated as well as the skin territory with an overlapping innervation (brown). Blue dots indicate single unit receptive field centers (compiled data from the current study, data recorded earlier(Milenkovic et al., 2014) and unpublished experiments). Digits (D), interdigital (I), thenar (T) and hypothenar (H) pads are indicated *Right:* Outside out configuration used in electrophysiological experiments (illustrated as mirror images). Dotted lines indicate the receptive fields of **C** the saphenous nerve, **F** the lateral and medial plantar nerve and I the median and ulnar nerve.

The receptive fields of myelinated axons were localized throughout the saphenous nerve territory (Fig. 1B), no receptive fields of saphenous nerve afferents were found to innervate adjacent glabrous skin (Fig. 1A-C). In our previous study we chiefly recorded from thinly myelinated polymodal nociceptors with axons in the tibial nerve innervating the hind paw glabrous skin (Milenkovic et al., 2014). Here we focused our analysis on afferents with myelinated axons that project to the glabrous skin via the lateral and medial plantar nerves that can be seen in the inside-out version of the *ex vivo* preparation (Fig. 1D). Receptive fields were found throughout the hind paw glabrous skin, but axons within the tibial nerve that project to the digits at the hairy/glabrous transition zone also sometimes innervated hairs (Fig. 1D-F). We routinely cut the sural nerve that contains some axons that innervate a thin sliver of glabrous skin on the lateral edge of the foot (Smith et al., 2013).

We developed a novel preparation that allowed us to make recordings from myelinated sensory afferents innervating the forepaw glabrous skin. The forepaw glabrous skin is innervated by the median and ulnar nerves. Recordings from skin regions with afferents in either nerve revealed that the functional nerve territories show some overlap in the middle of the paw. For example, digit 4 was innervated by afferents from both the median and ulnar nerves. As in the hind paw, the majority of axons within the median and ulnar nerves formed a receptive field within the glabrous skin, but hairy skin at the border of the glabrous skin (e.g. at the wrist or digits) was occasionally innervated by these axons (Fig. 1G-I). We noted that it was extremely important to remove muscle tissue surrounding the nerve branches in both the forepaw and hind paw glabrous skin preparations in order to maintain tissue viability. In the best cases recordings could be made for up to 8 hours following tissue removal. Note that all recordings in this study were made with the skin in the outside-out configuration (illustrated in Fig. 1 C,F,I), thus stimuli were delivered to the skin surface as would be the case *in vivo*.

### Receptor properties of Aβ-fiber and Aδ-fiber afferents across skin regions

Each single receptor-unit could be classified according to conduction velocity and stimulus response properties as one of four types of mechanoreceptor (Table 1). To apply quantitative mechanical stimuli to the receptive field we used a Piezo-driven actuator equipped with a force measurement device (Fig. 2A). This set-up allowed us to apply vibration stimuli of different frequencies with increasing amplitude, a protocol that allows the determination of force threshold for activation of mechanoreceptors. An example of such a determination for a rapidly-adapting mechanoreceptor (RAM) that responds to a 20Hz vibration is shown in Fig. 2B. We also used ramp and hold stimuli in which the 2 second long static phase was supra-threshold to activate the receptor and the velocity of the ramp was varied to probe the dynamic (or velocity) sensitivity of the receptor (Fig. 2C). Analysis of spike rates only during the ramp phase of the stimuli was used to assess the velocity sensitivity of the receptor. The example shown in Fig. 2C is from a slowly-adapting mechanoreceptor (SAM) which fires at much higher rates during the dynamic phase of the stimulus compared to the static phase. This stimulus protocol was applied to all afferent types which show dynamic responses to moving stimuli, i.e. RAMs, SAMs and D-hair receptors (Heidenreich et al., 2012; Lechner and Lewin, 2013; Zimmerman et al., 2014). Most nociceptors do not respond to skin movement but rather code static intensity. In this study we recorded the mechanosensitivity of SAMs and Aδ-fibers with nociceptor properties (so called Aδ-mechanonociceptors, AMs) across different skin regions. The receptor properties of these fibers were probed with a series of ramp and hold stimuli of constant velocity in which the amplitude was increased incrementally from 15-300mN (Fig. 2D).

**Figure 2:**
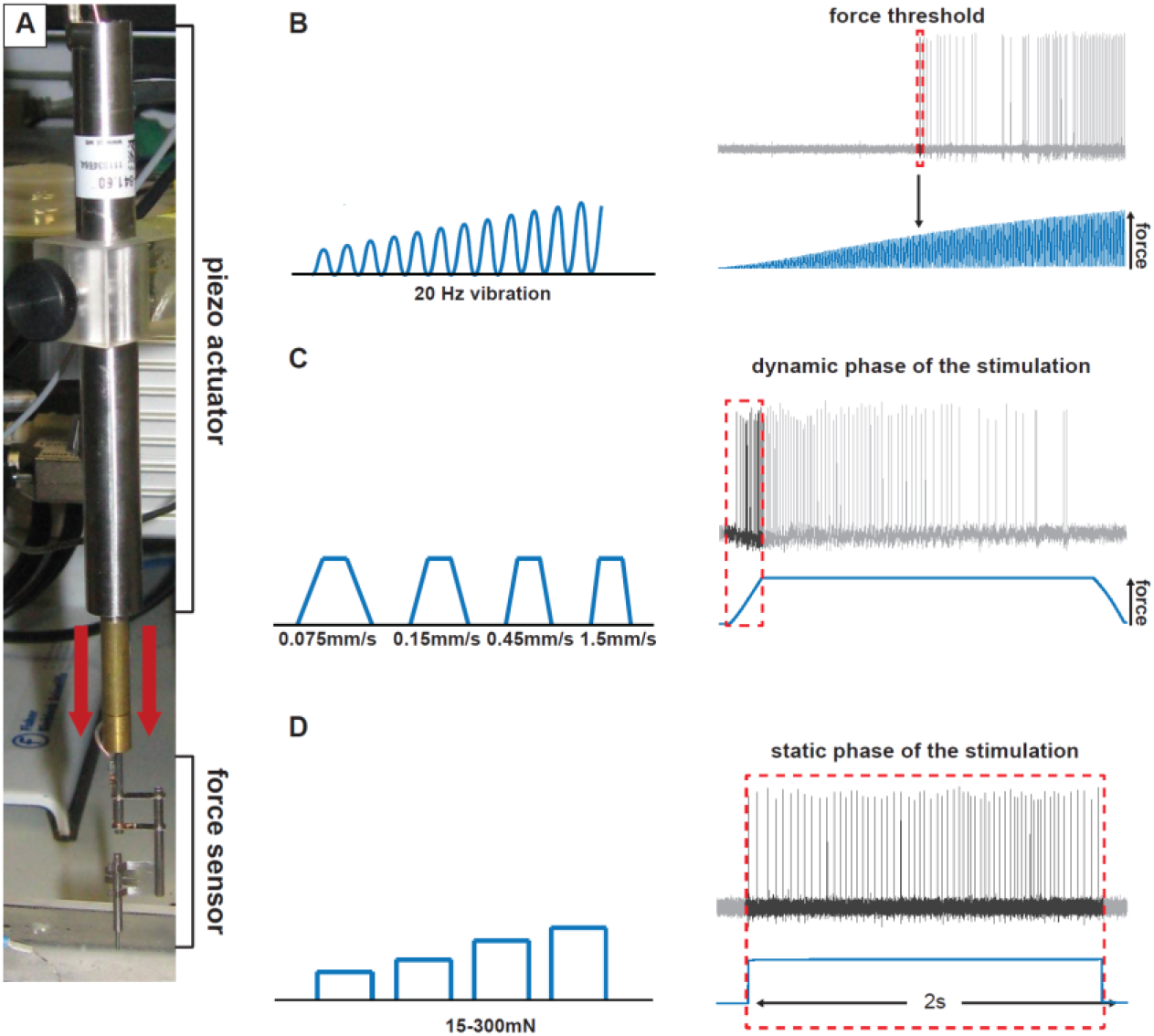
Stimulation protocols. **A** Example picture of the stimulation motor (piezo actuator, Physik Instrumente) and the force feedback system (force sensor, Kleindiek Nanotechnik). **B** *Left:* Schematic illustration of the sigmoidal vibrating stimulus (20Hz) with increasing intensity. *Right:* Example trace of a receptor fiber responding to the vibrating stimulus. The force at the time of the first action potential was measured. **C** *Left:* schematic illustration of the ramp and hold stimulation. Four different velocities were used *Right:* Example trace of a receptor fiber responding to a ramp and hold stimulation. Only the spikes at the dynamic phase of the stimulation were measured. **D** *Left:* Schematic illustration of the ramp and hold stimulation, four different intensities were used *Right:* Example trace of a receptor fiber responding to a ramp and hold stimulation. Only the spikes during the static phase of the stimulation were quantified.

**Table 1.**
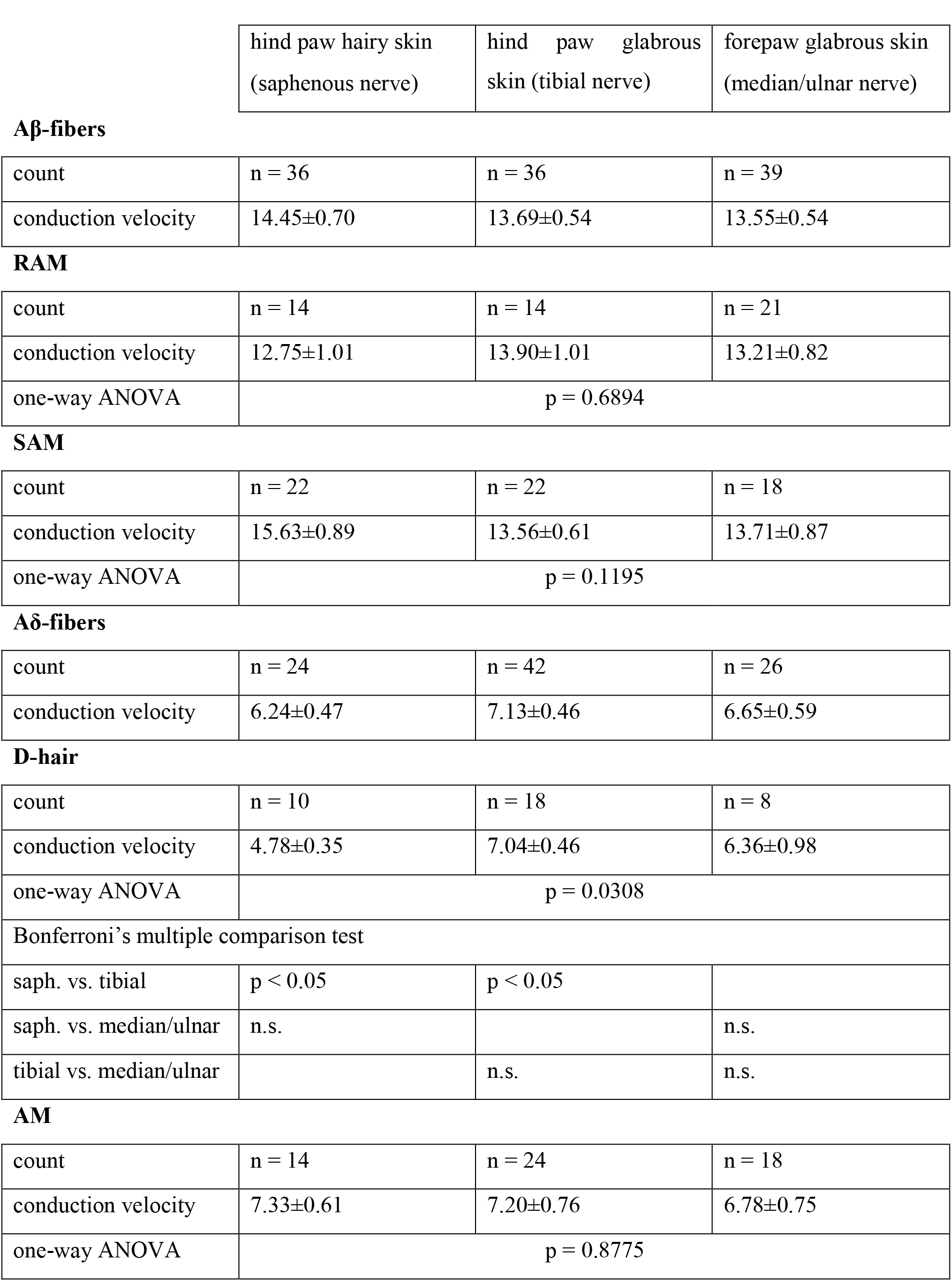

| hind paw hairy skin<br>(saphenous nerve) | hind paw glabrous<br>skin (tibial nerve) | forepaw glabrous skin<br>(median/ulnar nerve) |
| --- | --- | --- |

**Aβ-fibers**
|  |  |  |  |
| --- | --- | --- | --- |
| count | n = 36 | n = 36 | n = 39 |
| conduction velocity | 14.45±0.70 | 13.69±0.54 | 13.55±0.54 |

**RAM**
|  |  |  |  |
| --- | --- | --- | --- |
| count | n = 14 | n = 14 | n = 21 |
| conduction velocity | 12.75±1.01 | 13.90±1.01 | 13.21±0.82 |
| one-way ANOVA | p = 0.6894 |  |  |

**SAM**
|  |  |  |  |
| --- | --- | --- | --- |
| count | n = 22 | n = 22 | n = 18 |
| conduction velocity | 15.63±0.89 | 13.56±0.61 | 13.71±0.87 |
| one-way ANOVA | p = 0.1195 |  |  |

**Aδ-fibers**
|  |  |  |  |
| --- | --- | --- | --- |
| count | n = 24 | n = 42 | n = 26 |
| conduction velocity | 6.24±0.47 | 7.13±0.46 | 6.65±0.59 |

**D-hair**
|  |  |  |  |
| --- | --- | --- | --- |
| count | n = 10 | n = 18 | n = 8 |
| conduction velocity | 4.78±0.35 | 7.04±0.46 | 6.36±0.98 |
| one-way ANOVA | p = 0.0308 |  |  |
| Bonferroni's multiple comparison test |  |  |  |
| saph. vs. tibial | p < 0.05 | p < 0.05 |  |
| saph. vs. median/ulnar | n.s. |  | n.s. |
| tibial vs. median/ulnar |  | n.s. | n.s. |

**AM**
|  |  |  |  |
| --- | --- | --- | --- |
| count | n = 14 | n = 24 | n = 18 |
| conduction velocity | 7.33±0.61 | 7.20±0.76 | 6.78±0.75 |
| one-way ANOVA | p = 0.8775 |  |  |

### Rapidly-adapting mechanoreceptors (RAMs)

RAMs respond exclusively to moving stimuli, rapidly ceasing to fire after cessation of movement (Fig. 3A). Most RAMs in the hairy skin are associated with hair follicles, while the RAMs of the glabrous skin are associated with Meissner’s corpuscles (Heidenreich et al., 2012; Lechner and Lewin, 2013; Poole et al., 2014a; Zimmerman et al., 2014). Forty-nine of the 111 Aβ-fibers recorded could be classified as RAMs and these receptors were equally sampled across hind paw hairy skin, hind paw glabrous skin and forepaw skin (Table 1). We used increasing amplitude vibration stimuli to assess the mean force threshold to activate all RAMs that innervate the different skin regions. The mean force thresholds were, as expected, very low with thresholds <3mN. There was a tendency for forepaw afferents (median and ulnar nerve) to be activated with smaller forces compared to afferents in hairy skin (saphenous nerve) or those innervating the hind paw glabrous skin (tibial nerve), but this difference was not statistically significant (Fig. 3B; saphenous: n = 13; tibial: n = 13; median/ulnar: n = 21 Kruskal-Wallis test p = 0.6476).

**Figure 3:**
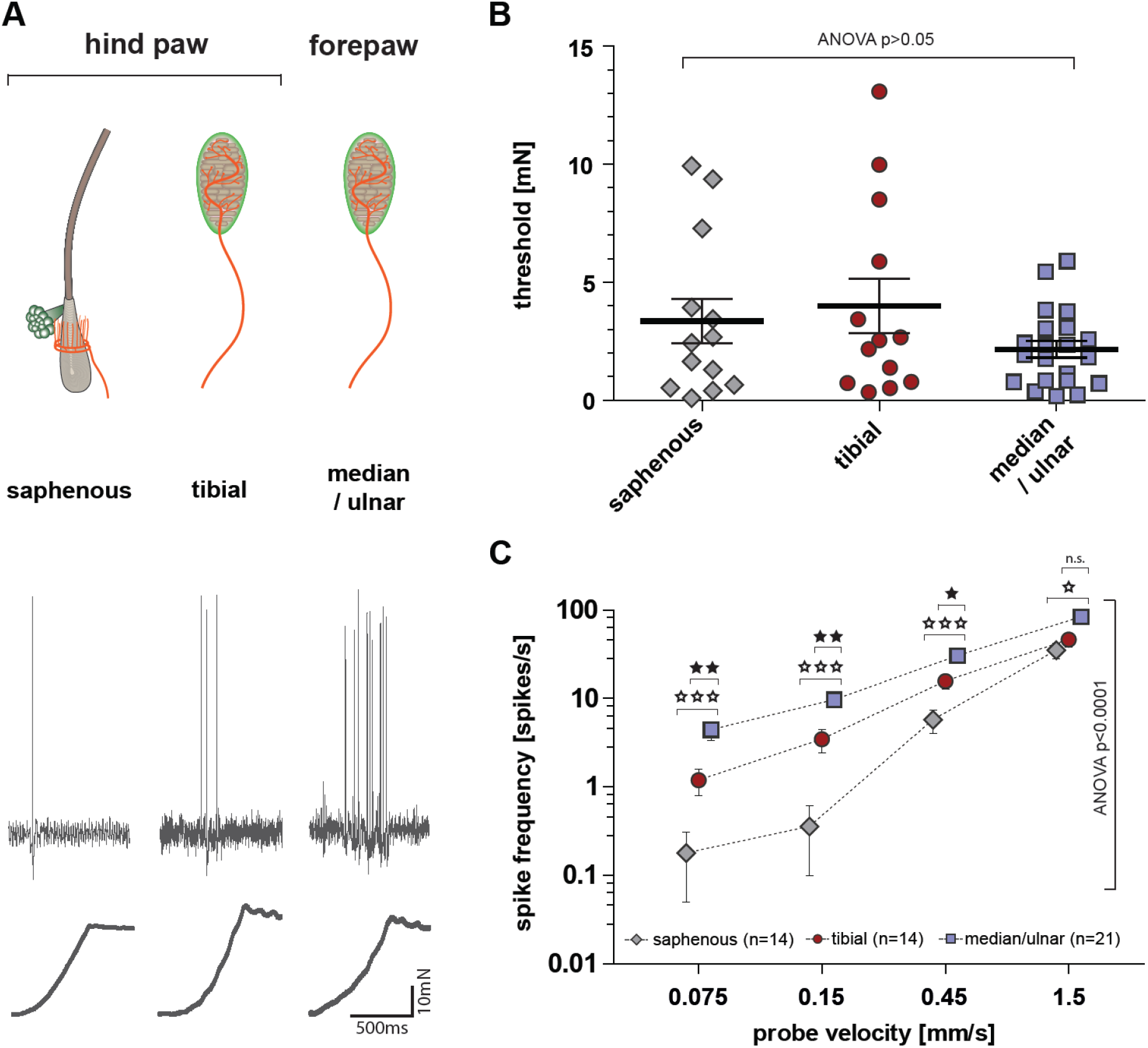
Response properties of RAM-receptors. **A** *top:* Schematic drawing of the predominant RAM anatomy in saphenous-nerve preparation (hair follicle receptor), tibial-nerve preparation (Meissner’s corpuscle) and median/ulnar-nerve preparation (Meissner’s corpuscle) *Bottom:* representative example traces of RAMs in response to a ramp and hold stimulation with a velocity of 0.45mm/s. **B** Minimal stimulation force needed to evoke an action potential in response to increasing amplitude vibrating stimuli (20 Hz); ANOVA: p>0.05; error bars represent SEM. **C** Average spike frequency in response to moving stimuli. Repeated measures ANOVA: p<0.0001; Bonferroni post-hoc tests are indicated; *** = p<0.001 ** p<0.01 * p<0.05; error bars represent SEM.

There was a quantitatively large difference in the way that RAMs in the glabrous forepaw skin coded moving stimuli compared to RAMs innervating hairy skin (Fig. 3C). Forepaw RAMs responded reliably with high firing rates to the slowest ramps used (0.075mm/s and 0.15mm/s), whereas hairy skin RAMs barely responded to the same stimuli (Fig. 3A,C) and these differences were statistically significant, (saphenous: n = 14; tibial: n = 14; median /ulnar: n = 21 repeated measure ANOVA F(2, 46) = 15.22, p< 0.0001). Thus RAMs in the forepaw glabrous skin are tuned to slower movements, but also have firing rates that were several fold higher than observed for afferents innervating hairy skin (Fig 3A,C); 0.075mm/s (saphenous vs. median/ulnar) p <0.001; 0.15mm/s (saphenous vs. median/ulnar) p<0.001; 0.45mm/s (saphenous vs. median/ ulnar) ANOVA with Bonferroni post-hoc test p <0.001; 1.5mm/s (saphenous vs. median/ulnar) p<0.05. The vast majority of the forepaw RAMs had receptive fields clearly located in the glabrous skin and therefore likely innervate Meissner’s corpuscles. It is possible that a very small number of receptors innervated hair follicles on the glabrous/hairy skin border, but due to the receptive field distributions (Fig. 1) these afferents could not make up more than 10% of the total. The majority of RAMs in the tibial nerve also innervate glabrous skin and therefore also likely have Meissner’s corpuscle endings. However, the sensitivity of RAMs found in the glabrous hind paw skin did not match that of the forepaw RAMs (Fig. 3A,C; ANOVA with Bonferroni post-hoc test: tibial vs. median/ulnar 0.075mm/s p <0.01; 0.15mm/s p<0.01; 0.45mm/s p <0.05; 1.5mm/s p>0.05). Glabrous hind paw RAMs were, however, capable of coding slower velocities than RAMs in hairy skin, but this was not statistically significant (Fig. 3C).

Because presumptive Meissner’s corpuscle receptors located in the glabrous forepaw were more responsive to ramp stimuli than those in the hind paw, we thus questioned whether receptor morphology differed between these two locations. In the first step we investigated the number of nerve fibers associated with each Meissner’s corpuscle. We used immunofluorescence methods to visualize Meissner’s corpuscles and their innervation in hind paw and forepaw glabrous skin. We used antibodies against S100 which is found in terminal Schwann cells that form the corpuscle and antibodies against NF200 to visualize the myelinated axons that innervate the corpuscle (Fig 4A,B). We examined a total of 175 Meissner’s corpuscles (84 located in the hind paw glabrous skin and 91 located in the forepaw glabrous skin) and counted the number of myelinated nerve fibers innervating each corpuscle (Fig. 4D). Evidently each corpuscle in the forepaw skin was on average innervated by more axons than those in the hind paw. Thus, more than 50% of corpuscles were found with 2-3 innervating axons in the forepaw whereas in the hind paw 70% of the corpuscles were innervated by 1-2 axons only (Fig. 4C; hind paw: median = 2, mean = 2.02 ± 0.09 fibers, n = 84; forepaw: median = 3, mean = 2.57 ± 0.91 fibers, n = 91; mean ± SEM; Mann-Whitney U-test, p <0.0001). It could be that the increased innervation of Meissner’s corpuscles in the forepaw is simply due to larger corpuscles in this skin area. However, we measured the volume (V) of 135 Meissner’s corpuscles (65 from hind paw and 70 from forepaw skin) and found that there was no difference in their mean volumes (Fig 4E; hind paw: V = 3.53 ×10 μm^3^, n = 65; forepaw: V = 3.52 ×10^3^ μm^3^, n = 70; mean ± SEM; Mann-Whitney U-test, p=0.6645). We next measured the overall density of Meissner’s corpuscles in the forepaw and hind paw glabrous skin. Using confocal microscopy we could image a large volume of skin and measure precisely the number of corpuscles in a known volume (see Supplementary Movie 1). We chose to image the elevated interdigital pads of the glabrous paw skin, sometimes called “running pads” (indicated in Fig. 1E,F,H,I), as these are discrete and easily identifiable regions that can be directly compared between fore and hind paw skin. We measured the average number of Meissner’s corpuscles per volume [μm^3^] of skin (Fig. 4B, top) within the interdigital pads III closest to the second digit of the forepaw or the corresponding interdigital pad IV of the hind paw (n=7 animals, age = P23-28). On average 34.6 ± 2.7 Meissner’s corpuscle in the forepaw and 43.3 ± 3.4 Meissner’s corpuscles in the hind paw were counted within one interdigital pad. Thus, the average density of Meissner’s corpuscles was significantly higher in the forepaw glabrous skin interdigital pad III with 1.10 ± 0.09 corpuscle/10^−6^μm^3^ compared to interdigital pad IV of the hind paw with 0.81 ± 0.03 corpuscle/10^−6^μm^3^ (Fig. 4E; paired t-test: p = 0.0188).

**Figure 4:**
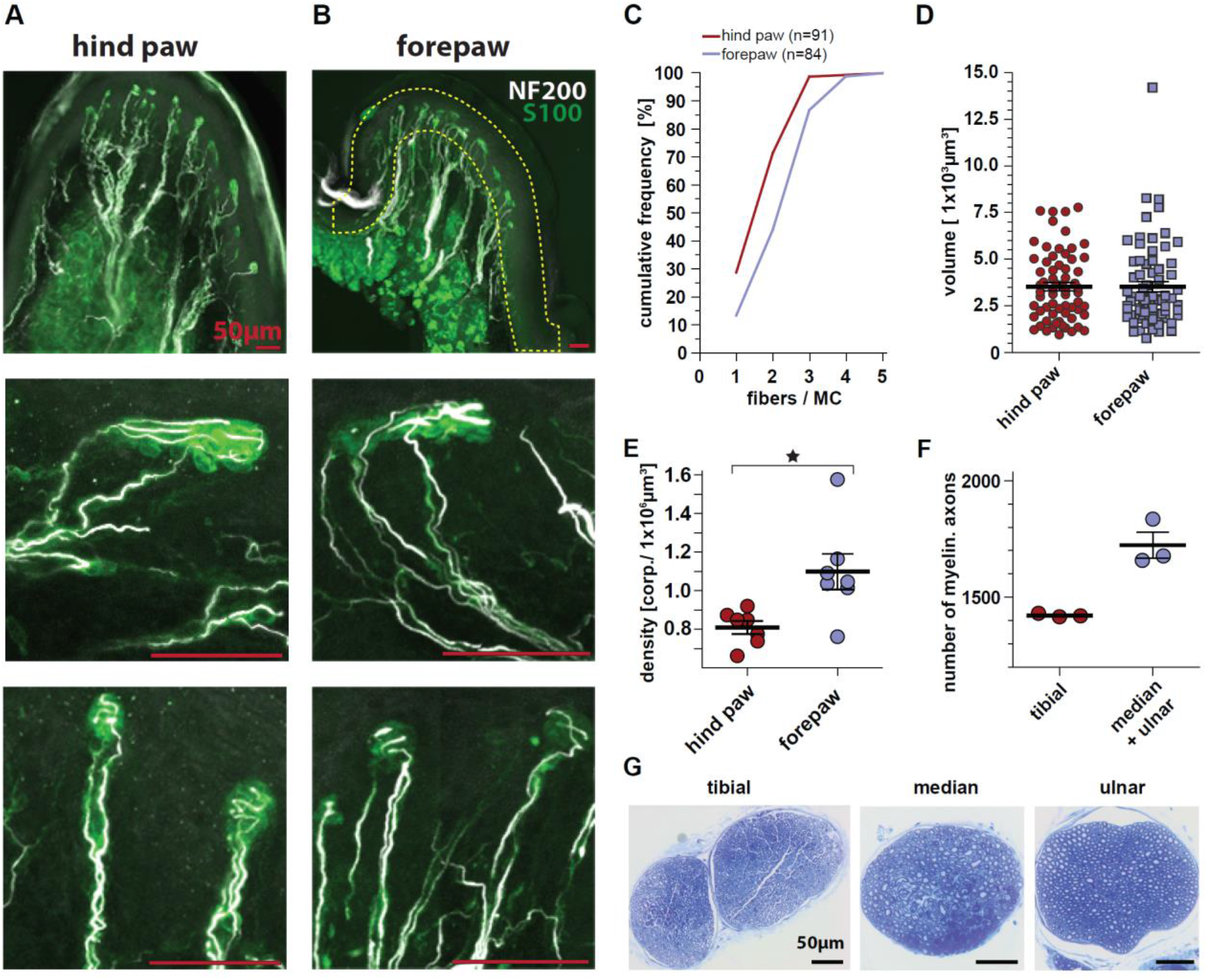
Meissner’s corpuscle anatomy. Sagittal vibratome sections of **A** the interdigital pad IV of the hind paw and **B** of the interdigital pad III of the forepaw. **A** and **B** *top:* Immunofluorescence picture of a running pads with labeled Meissner’s corpuscle (anti S100) and myelinated nerve fibers (anti-NF200) *middle* and *bottom:* Magnified representation of single Meissner’s corpuscles. **C** Cumulative frequency plot of the number of fibers innervating a single Meissner’s corpuscle. Lower and upper quartile as well as the median are indicated. **D** Size (volume) of Meissner’s corpuscle in the hind and forepaw running pads; error bars represent SEM. **E** Meissner’s corpuscle density in the interdigital running pads III (hind paw) and IV (forepaw); paired t-test: p= 0.0188; error bars represent SEM. **F** Number of myelinated axons counted in the tibial and the median plus the ulnar nerve; error bars represent SEM. **G** Representative semi-thin microscopy pictures used to quantify myelinated axon number. All scale bars in panels **A** and **B** are 50 μm

We used semi-thin sections of peripheral nerves to make quantitative estimates of the total number of myelinated fibers in the tibial nerve that innervate the hind paw glabrous skin compared to the number of myelinated fibers in median and ulnar nerves that innervate the forepaw glabrous skin. The mean myelinated axon count for the median and ulnar nerves combined was 1723 ± 56 (n = 3) compared to 1422 ± 4 (n = 3) for the entire tibial nerve (Fig 4F). We estimated the innervated skin area of the hind paw glabrous innervated by the tibial nerve to be 2.9 larger than the innervated area of the forepaw glabrous skin innervated by the median and ulnar nerves (median/ulnar: 65 ± 2.2 fibers/mm^2^; tibial: 22 ± 0.4 fibers/mm^2^, 4 animals age P23-28; mean ± SEM). Therefore, it is obvious that the forepaw glabrous skin has as much as a fourfold higher afferent innervation density in terms of myelinated fibers compared to the hind paw glabrous skin.

### Slowly-adapting mechanoreceptors (SAMs)

Slowly-adapting type mechanoreceptors respond well to both indentation and vibration stimuli. To compare SAMs in hairy skin to those with receptive fields in glabrous skin, a ramp and hold stimulus was used. The response of a typical SAM is shown in Fig. 5A. Of the 111 Aβ-fibers recorded across all skin areas 62 were SAMs: 22 from the hind paw hairy skin, 22 from the hind paw glabrous skin and 18 from the forepaw glabrous skin. We calculated the mean firing rate of all SAMs to supra-threshold ramp and hold stimuli (2 seconds long) over time by binning spike counts into 100 ms bins (Fig. 5B). As expected, mean firing rates were highest during the dynamic phase (up to 150Hz) and adapted to lower rates during the static phase (10-30Hz). Importantly, no significant differences in the spike rates were found for SAMs recorded across the three skin regions (Fig. 5B saphenous: n = 20; tibial: n = 20; median/ulnar: n = 18) repeated measure ANOVA F(2, 55) = 1.20; p = 0.3087). We also used a series of increasing ramp velocities with constant supra-threshold force to probe SAM velocity sensitivity as described above for RAMs (Fig. 5C). The stimulus response functions were typical for SAMs and the velocity sensitivities of the receptors were almost identical across the three skin regions tested (Fig 5C saphenous: n = 21; tibial: n = 22; median/ulnar: n = 16 repeated measures ANOVA F(2, 56) = 1.11; p = 0.3364). We next tested the sensitivity of the SAMs to static indentation using a series of stimuli varying from 15 up to 250mN. The mean rates of firing reached a plateau around 60mN for SAM receptors across all three skin regions confirming that these receptors are tuned to detect low intensity threshold static indentation (Fig 5D). Across the saphenous nerve (hind paw hairy skin), the tibial nerve (hind paw glabrous skin) and the median/ulnar nerve (forepaw glabrous skin) there was no statistically significant difference in the mean stimulus response for the static component between SAMs (Fig 5D saphenous: n =19; tibial: n = 16; medial/ulnar: n =16, repeated measures ANOVA F(2, 48) = 0.23; p = 0.7923). Finally, we estimated the force threshold for SAMs using a 20Hz vibration stimulus. As expected, most SAMs had very low thresholds for activation (mostly between 1-5mN) and the mean thresholds were not statistically significantly different from each other across the three skin regions (Fig 5E saphenous: n = 22; tibial: n = 19; median/ulnar: n = 17 Kruskal-Wallis test p = 0.6444). By analyzing the variance of firing rates during static indentation it is possible to distinguish between SAM type I receptors innervating Merkel cells and SAM type II receptors that do not innervate Merkel cells (Wellnitz et al., 2010). Analyzing our data set using this method revealed no differences in the proportion of SAM type I and SAM type II receptors across skin regions. In addition, the receptor properties of both SAM types were not different across the three skin regions (data not shown).

**Figure 5:**
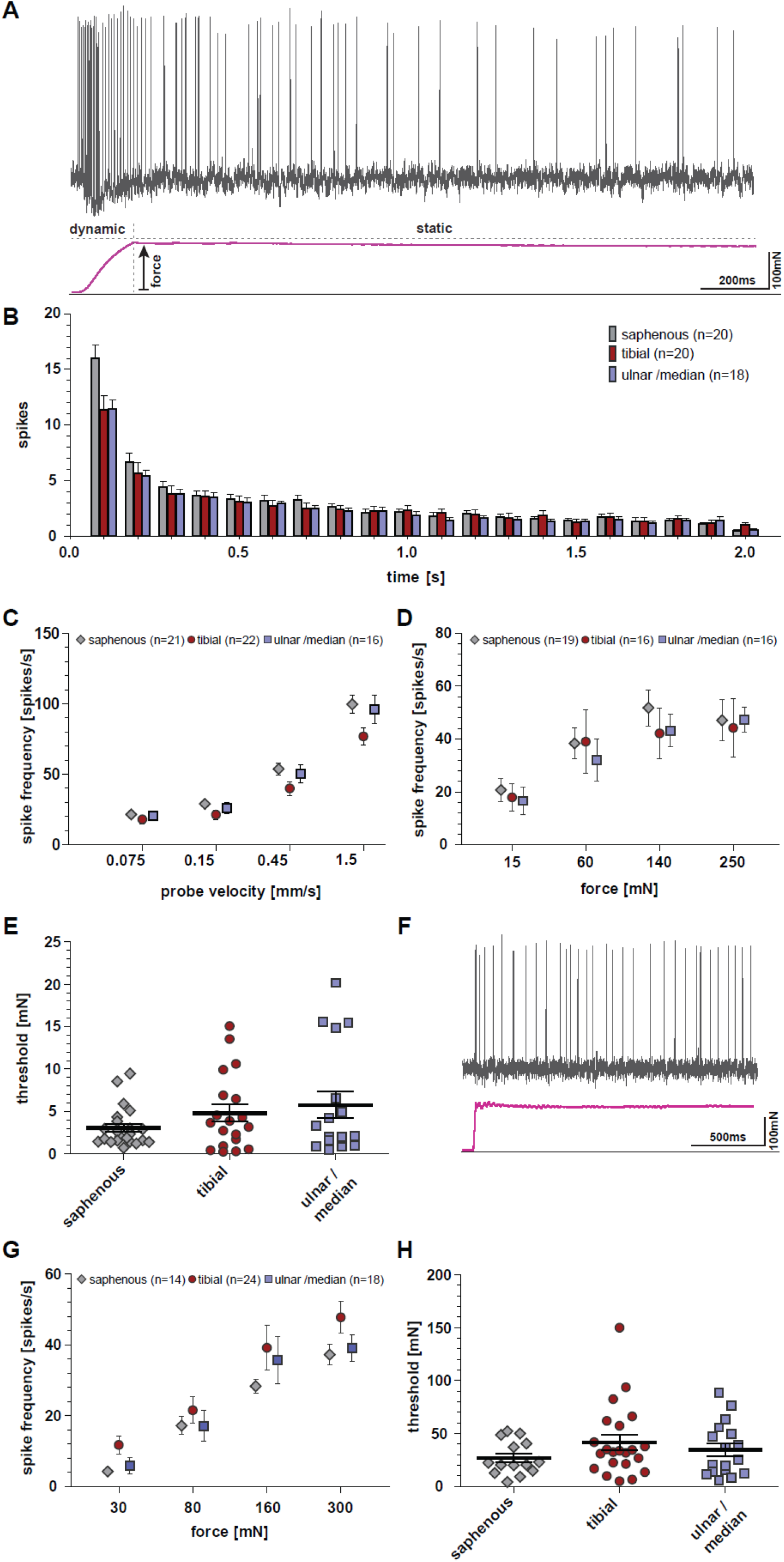
Receptor properties of SAM. **A** Representative example trace of a SAM response (hind paw glabrous skin) to a ramp and hold stimulation. **B** Average spike count (bin 0.1s) to a suprathreshold mechanical stimulus over 2 second duration. **C** Average spike frequency of SAMs in response to moving stimuli; repeated measure ANOVA: p > 0.05; error bars represent SEM. **D** Average spike frequency of SAMs in response to ramp and hold stimulation; repeated measure ANOVA: p > 0.05; error bars represent SEM. **E** Minimal force needed to evoke an action potential of SAMs in response to a 20 Hz vibrating stimuli; repeated measure ANOVA: p > 0.05; error bars represent SEM. **F** Representative example trace of a AM response (hind paw glabrous skin) to a ramp and hold stimulation. **G** Average spike frequency of AMs in response to ramp and hold stimulations; repeated measure ANOVA: p > 0.05; error bars represent SEM. **H** Minimal force needed to evoke an action potential in AMs in response to a fast moving ramp stimulation; repeated measure ANOVA: p > 0.05; error bars represent SEM.

### A-δ fiber mechanonociceptors (AMs)

Myelinated nociceptors, also called A-mechanonociceptors (Lewin and Moshourab, 2004), are thought not to be associated with specific structures or corpuscles in the skin (Arcourt et al., 2017; Kruger et al., 1981). Myelinated nociceptors have been extensively studied in the hairy skin and typically respond with increasing firing rates to increasing static force (Garell et al., 1996; Lewin and Moshourab, 2004; Milenkovic et al., 2008). A total of 56 Aδ-mechanonociceptors were studied; 14 with a receptive field in hind paw hairy skin, 24 with a receptive field in hind paw glabrous skin and 18 with receptive fields in forepaw glabrous skin. Stimulus response functions to steadily increasing ramp and hold stimuli with intensities ranging from 30mN to 300mN were plotted (Fig. 5F,G). The stimulus response functions of AMs found in all three skin areas were indistinguishable from each other and there was no statistically significant difference between them (Fig. 5G), repeated measures ANOVA; saphenous: n = 14; tibial: n = 24; median/ulnar: n = 18, F(2, 53) = 2.15, p = 0.1262. For each AM unit we also measured the minimum force necessary to trigger the first action potential and again noted that in each skin area comparable high mechanical thresholds were found (Fig. 5H). Mean thresholds were slightly higher in the two glabrous skin areas, but this difference was not significantly different (Kruskal-Wallis test (saphenous: n = 14; tibial: n = 22; median/ulnar: n = 17) p = 0.4887). Thus, the mechanosensitive properties of A-mechanonociceptors appear to be uniform across different skin areas.

### D-hair receptors

D-hairs are the most sensitive mechanoreceptors in the skin and generally have the largest receptive fields (Burgess et al., 1968; Koltzenburg et al., 1997; Lechner and Lewin, 2013; Leem et al., 1993; Lewin and McMahon, 1991; Wang and Lewin, 2011). A total of 36 D-hair receptors were recorded: 10 from the hind paw hairy skin (saphenous nerve), 18 from the hind paw glabrous skin (tibial nerve) and 8 from the forepaw glabrous skin (median and ulnar nerves) (Table 1). All recorded D-hair receptors had conduction velocities in the Aδ-fiber range (between 1-10m/s), however, the mean conduction velocities of D-hair receptors in nerves innervating glabrous skin (tibial, median and ulnar nerves) were significantly faster than those recorded in the saphenous nerve which exclusively innervates hairy skin (Table 1). We used the same protocols to test velocity sensitivity of D-hair receptors as we had used for RAMs. D-hair receptors are especially sensitive to fast moving stimuli (Fig. 6A,B) and had very low mechanical thresholds to vibration stimuli (Fig. 6C). There was no significant difference in the mean mechanical thresholds of D-hair receptors recorded across the three skin areas (Fig 6C saphenous: n = 13; tibial: n = 10; median/ulnar: n = 5, Kruskal-Wallis test p = 0.41). Importantly, the coding properties of all D-hair receptors recorded were almost identical across all three skin areas (Fig. 6B,C saphenous: n = 10; tibial: n = 18; median/ulnar: n = 7; repeated measures ANOVA F(2, 32) = 1.50; p = 0.24).

**Figure 6.**
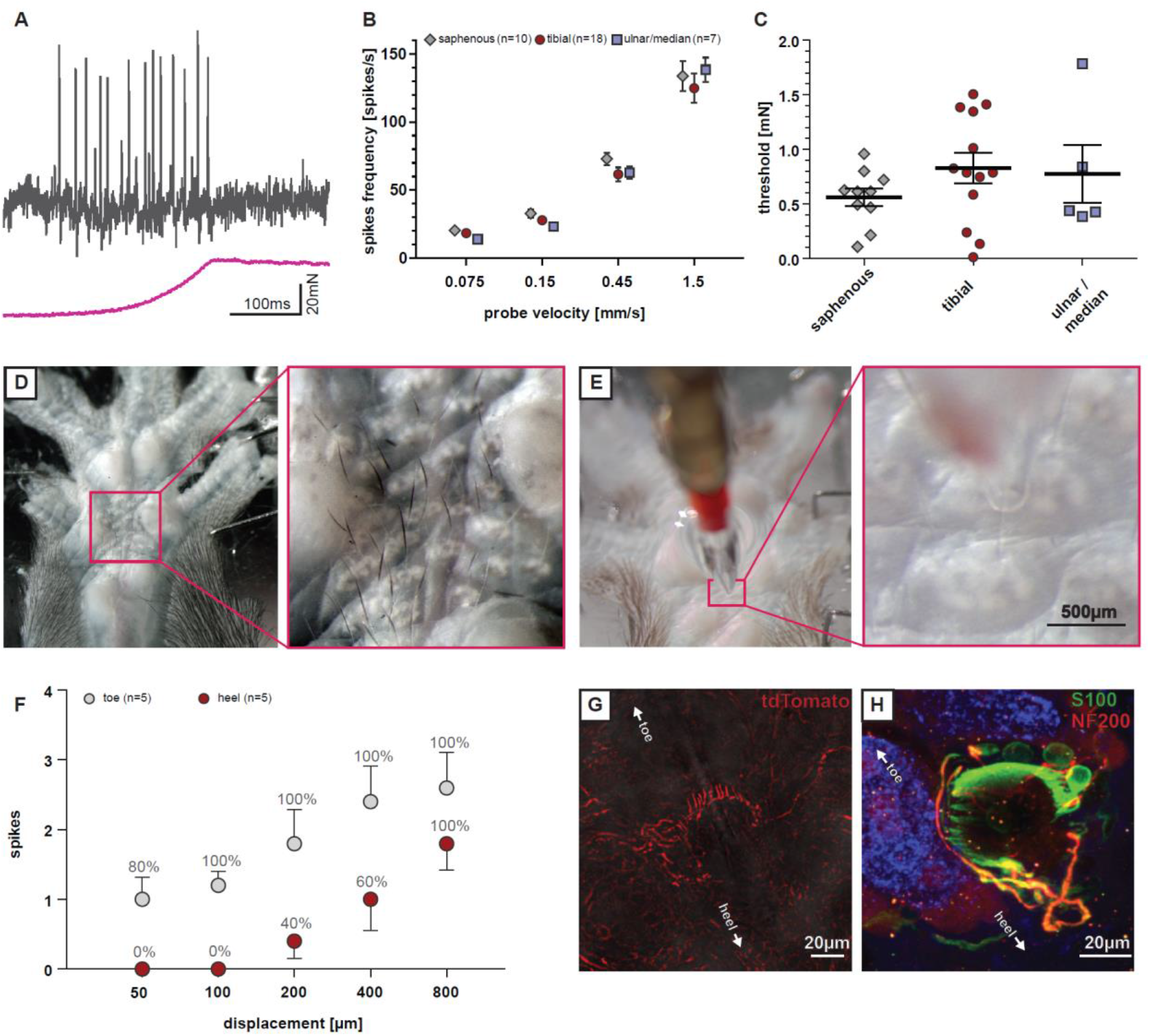
Functional and anatomical properties of glabrous skin D-Hair receptors. **A** Representative example traces from D-hair receptors recordings in response to a ramp and hold stimulation with a velocity of 0.45mm/s. **B** Average spike frequency in response to moving stimuli. Repeated measure ANOVA: p>0.05; error bars represent SEM. **C** Minimal stimulation force needed to evoke an action potential in response to increasing vibrating stimuli (20 Hz); ANOVA: p>0.05; error bars represent SEM. **D** Hairs at the glabrous hind paw skin; right panel: magnification **E** Experimental setup of the hair deflection experiment: The mounted hair can be deflected in the direction towards the toe, the heel or sideways; right panel: magnification to display the hair partially inside the glass capillary. **F** D-hair receptors respond with more action potentials (average spikes) and respond more reliable (% of receptors responding) to small hair deflections in the direction towards the toe compared to the direction of the heel. Error bars represent SEM. **G** Ca_v_3.2 positive nerve fibers cluster at one side of the follicle (whole mount preparation) **H** Whole mount hair follicle receptor staining; TSC in green (anti-S100), myelinated nerve fibers in red (anti-NF200) and auto fluorescence in the blue channel.

We recorded many D-hair receptors with axons in the tibial nerve; some of these afferents had receptive fields in the hairy skin at the transition zones between glabrous and hairy skin at the heel and toes. However, we discovered a group of very fine hairs (~20-30 in total) in the central region of the glabrous skin surrounded by the so-called running pads in all mice examined (Fig. 6D). Around half of the D-hair receptors recorded from the tibial nerve had a receptive field in this area and were specifically activated by movements of these very fine hairs. We rarely found classical RAMs that could be clearly activated by movement of the same set of hairs. When we observed such units it was impossible to determine if the movement of the hair was actually activating Meissner receptor endings in the near vicinity. These observations led us to hypothesize that these glabrous hind paw hairs are almost exclusively innervated by D-hair receptors.

Recently, it was shown that D-hair receptors in the back skin show direction sensitivity (Rutlin et al., 2014). The hairs in the middle of the hind paw glabrous skin are unusual in that the space between individual hairs is so large that it is straightforward to move a single hair without any danger of simultaneously manipulating adjacent hairs (Fig. 6E). We used a glass capillary to capture a single hair and by using a stepping motor we were able to move the hair in 2-dimensional space (see supplementary Video 2). As shown in Fig 5D, all these very fine hairs lie flat on the skin with hair growth orientated towards the toes. Placing the hair within the capillary meant that adjustment to a central position led to transient activation of the D-hair receptor, but firing ceased as soon as movement stopped. We then moved the hairs in the direction of growth, bending the hair in the direction of the toes, which led to the most robust activation of the receptor. In contrast, movement of the hair against its growth led to a weaker activation of the receptor. Thus, we were able to quantify the directional sensitivity of these receptors (Fig. 6F, Table 2, Supplementary Video 2). Movement of the hair from side to side generally led to an intermediate level of receptor activation (Table 2). The directional sensitivity of these specialized D-hair receptors means that their activation is greatest when the hairs are pushed against the skin. Therefore, under normal circumstances these receptors would be best activated as the animal places its foot on a surface. Activation would be least if the foot slides over a rough surface in the direction of forward movement.

**Table 1.** Responses of D-hairs to hair deflection

| Movement direction | Number of steps | 8 | 16 | 32 | 64 | 128 |
| --- | --- | --- | --- | --- | --- | --- |
| towards the toes | responding receptors | 4/5 | 5/5 | 5/5 | 5/5 | 5/5 |
|  | avg. number of spikes | 1 | 1.2 | 1.8 | 2.4 | 2.6 |
| towards the heel | responding receptors | 0/5 | 0/5 | 2/5 | 3/5 | 5/5 |
|  | avg. number of spikes | 0 | 0 | 0.4 | 1 | 1.8 |
| towards the sides<br>(average of both sides) | responding receptors | 1/5 | 1/5 | 2.5/5 | 5/5 | 4.5/5 |
|  | avg. number of spikes | 0.2 | 0.2 | 0.6 | 1.3 | 2.9 |

We took advantage of the fact that, in the adult, expression of the T-type calcium channel gene Ca_v_3.2 is highly specific for D-hair receptors (Shin et al., 2003; Sierra et al., 2017; Wang and Lewin, 2011). By using a Ca_v_3.2^cre^ mouse combined with adeno-associated virus transduction of sensory axons with AAV9 vectors carrying a floxed tdTomato reporter we could specifically label the peripheral terminals of sensory neurons that express the Ca_v_3.2 gene (Sierra et al., 2017). D-hair receptors form lanceolate endings around hair follicles which unlike RAMs do not form a closed circle around the follicle but instead display a horseshoe configuration in which one region of the follicle is devoid of endings (Rutlin et al., 2014; Sierra et al., 2017). We injected the adeno-associated AAV9-FLEX-tdTomato virus into the sciatic nerve of Ca_v_3.2 mice and harvested glabrous skin tissue from the same mice 8 weeks later. We found that in each of the four mice studied we could visualize lanceolate endings around single hair follicles in the glabrous skin (see Supplementary Video 3). D-hair lanceolate endings showed an asymmetrical innervation of the hair follicles so that the open part of the horseshoe was orientated toward the heel (Fig. 5G). In most preparations only a few follicles received a tdTomato-positive innervation consistent with the idea that the viral approach produces sparse labelling. We observed that single D-hair receptors were activated by movements of up to 9 of the hairs present, consistent with the idea that single afferents are labeled with the viral labelling approach. Using whole mount immunostaining in wild type mice we could visualize the innervation of these hair follicle receptors using primary antibodies against NF200, a marker for myelinated fibers, and S100, staining terminal Schwann cells (Fig. 6H). Immunofluorescence staining using S100 and NF200 antibodies confirmed that these hairs are innervated by myelinated hair follicle receptors that display lanceolate endings (Fig. 6H). In addition, we did observe some circumferential endings around these hairs (Fig. 6H).

### Glabrous D-hair receptors are evolutionarily ancient in rodents

The specialized hairs of the glabrous hind paw skin are likely exclusively innervated by D-hair receptors in laboratory mice. We never observed such hairs on the forepaw glabrous skin in mice. This raised the question as to whether such hairs, termed here glabrous D-hair receptors, may have arisen as a non-essential sensory trait through generations of in breeding in laboratory mice. We used C57Bl/6j mice in this study, but we also observed glabrous D-hair receptors in another inbred mouse strain CBA/J mice (Carter et al., 1952) (Fig. 7A,B).

**Figure 7.**
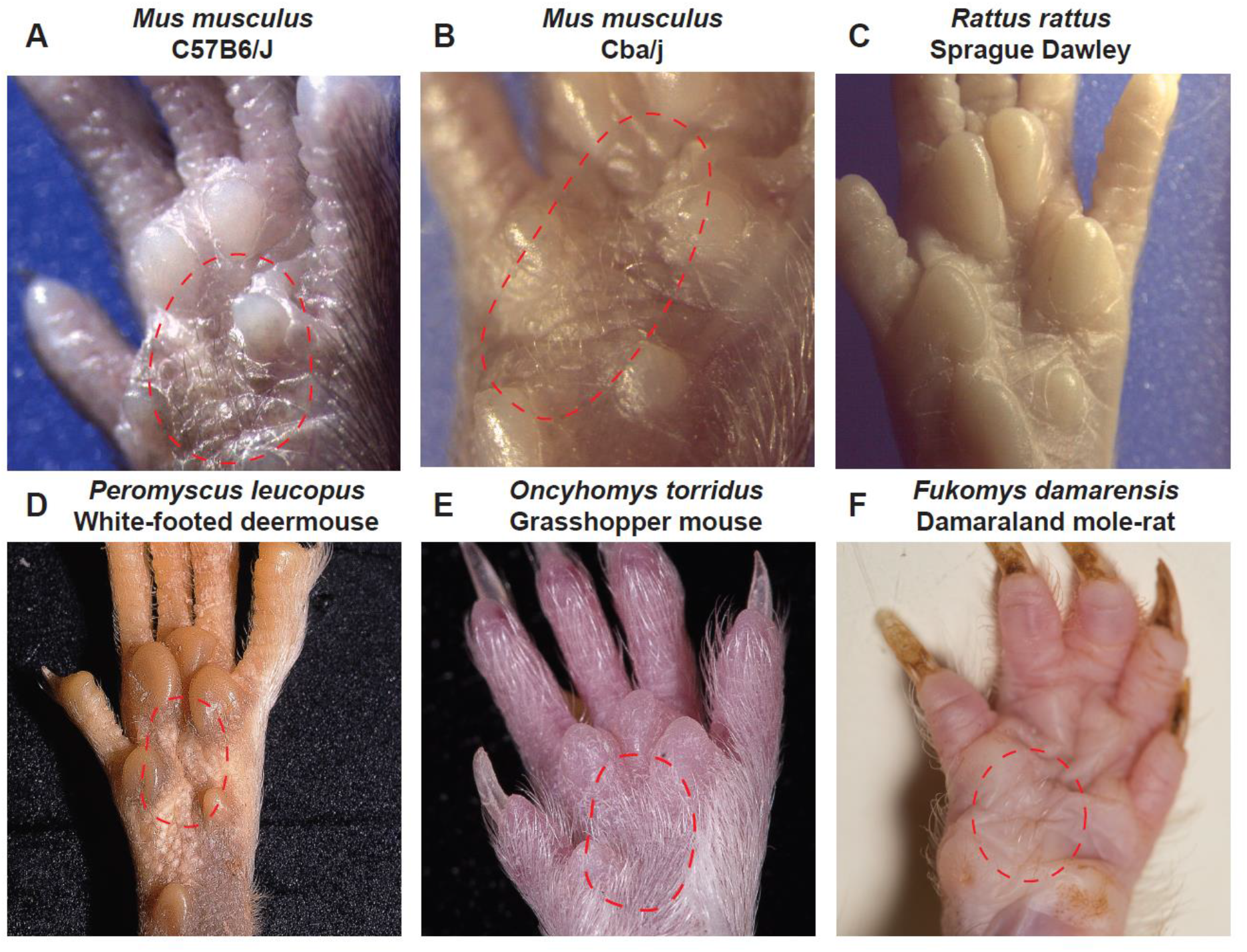
Glabrous skin D-Hair receptors are found in North American and African rodent species. **A – F** Representative pictures of the glabrous hind paw skin from laboratory rodents (top row) and three wild caught rodent species from Africa and North America. **A-B** Show the pictures of these very fine hairs in two laboratory in bred mouse strains **A** C57Bl6/J and **B** Cba/j mice. **C** We observed no hairs on the glabrous skin of laboratory rats. **D** Sparse fine hairs were observed on the hind paw glabrous skin of the North American white footed mouse and quite dense hairs were found on the same region in the Grasshopper mouse which has its habitat in the Arizona desert **E**. Very distinctive glabrous hairs were also observed on the glabrous skin of the Damaraland mole-rat **F**.

However, there was no evidence of glabrous D-hair receptors in inbred laboratory rats (*Rattus rattus*, Sprague Dawley strain) (Fig. 7C).

With over 2000 living species (~40% of all mammals) that populate most continents, rodents are highly successful and diverse (Don E. Wilson and DeeAnn M. Reeder, 2005). We reasoned that glabrous D-hairs may have been an ancient adaptation in this order that has appeared and reappeared during species diversification. We screened the hind feet of 3 North American rodents in the family Cricetidae and 9 African rodent species for the presence of suspected glabrous D-hair receptors. Two of the North American species, *Peromyscus leucopus* (white-footed deermouse) and *Oncyhomys torridus* (Southern Grasshopper mouse) had obvious small hairs within the same region of hind paw glabrous skin as seen in the mouse (Fig. 7D,E). The white-footed deermouse is distributed across the mid-western and eastern United States (Delaney and Hoekstra, 2018) whereas the Southern Grasshopper mouse is restricted to the deserts and grasslands of the western U.S. and northern Mexico. The white-footed deermouse is an opportunistic insectivore, while the southern grasshopper mouse unusually, is an obligate carnivorous rodent that feeds on insects and scorpions (Rowe and Rowe, 2015; Rowe et al., 2013). The third North American rodent species *Neotoma albigula* (white-throated woodrat) clearly lacked hind paw glabrous hairs (Supplementary Figure 1). This species is, like *Oncyhomys torridus*, also found in the dry woodlands and deserts of the southwestern U.S. and northern Mexico (Spencer and Spencer, 1941).

Among the African rodent species that we surveyed none of the surface dwelling species, like *Otomys sloggetti* (Ice rat), *Micaelamys namaquensis* (Namaqua rock mouse), *Saccostomys campestris* (South African Pouched mouse), and *Rhabdomys dilectus* (Four striped grass rat) possessed glabrous D-hair receptors (Supplementary Fig. 1). There is a rich diversity of exclusively underground living African mole-rat species from the family *Bathyergidae* found from the horn of Africa to the Cape region in Southern Africa. We surveyed the glabrous hind paws of 7 African mole-rats; *Heterocephalus glaber* (naked mole-rat), *Heliophobius emini* (Emin’s mole-rat), *Georychus capensis* (Cape mole-rat), *Bathyergus suillis* (Cape Dune mole-rat), *Cryptomys hottentotus mahali* (Mahali mole-rat), *Cryptomys hottentotus pretoriae* (Highveld mole-rat) and *Fukomys damarensis* (Damaraland mole-rat) with at least one representative species from each of the 6 genera of Bathyergidae examined. One of these species *Fukomys damarensis*, a eusocial species found in south western and central Africa (Bennett and Faulkes, 2000; Bennett and Jarvis, 1988; Davies et al., 2015), possessed discrete groups of glabrous D-hair receptors with remarkably similar morphology to those of the laboratory mouse (Fig. 7E).

## Discussion

Here we show that it is possible to record from mouse forepaw afferents using a novel *ex vivo* skin nerve preparation. By comparing the physiological properties of sensory afferents across three skin areas, we identified novel specializations of mouse forepaw glabrous skin receptors. Firstly, RAMs that innervate Meissner’s corpuscles are several fold more sensitive to slow skin movement compared to the same receptors in hind paw glabrous skin, or functionally similar RAMs in hairy skin. Second, individual Meissner’s corpuscles are innervated by ~25% more sensory axons than in other skin areas and are present at significantly higher densities in the forepaw compared to hind paw glabrous skin. Strikingly, all mechanoreceptors with myelinated axons are present at much higher density (>3 fold higher) in the forepaw skin compared to hind paw glabrous skin. Nevertheless, by recording from all sub-populations of myelinated mechanoreceptors as well as myelinated nociceptors we could show that it is only RAMs that show functional specialization in the forepaw. In addition, we also identified a unique population of D-hair receptors that provide an almost exclusive innervation to a small set of hairs within the hind paw glabrous skin of the mouse. These D-hair receptors show prominent directional sensitivity so that they are optimally activated when these very small hairs are pushed against the skin as would happen during a foot fall. Interestingly, these glabrous D-hair receptors are not ubiquitous in other rodent species, but we show that they have likely arisen very early in rodent evolution as we identified one African mole-rat species and two North American rodent species that appear to possess such hairs (from a total of 12 examined).

Our data suggests that the mouse has a highly developed forepaw tactile system adapted to manipulating objects and exploring their texture. The fact that myelinated sensory fiber receptor density was more than 3 fold higher than in the hind limb would enable much higher discriminative abilities using the forepaw. We calculated that the density of the myelinated sensory innervation was a striking 65 fibers/mm^2^. Analogously in humans tactile discrimination performance is clearly positively correlated with sensory innervation density which is highest on the finger tips and tongue (Van Boven and Johnson, 1994; Johansson and Vallbo, 1979a, 1979b). Indeed mechanoreceptor density on the finger tips has been estimated to be ~240/cm^2^ (Johansson and Vallbo, 1979c). We show here that Meissner’s corpuscle density is not only very much higher in forepaw compared to hind paw glabrous skin, but also that each corpuscle receives around 25% more sensory endings compared to an equivalent corpuscle in the hind paw (Fig. 4). The anatomical adaptations that we have described in forepaw Meissner’s corpuscles might underlie their functional specialization, including increased sensitivity to slower moving mechanical stimuli (Fig. 3). However, it is not clear why higher densities of sensory axons within a Meissner’s corpuscle should increase the velocity sensitivity of the sensory unit. Mechanoreceptors including RAMs that innervate Meissner’s corpuscles are thought to be equipped with a mechanotransduction apparatus that includes the mechanosensitive ion channel PIEZO2 and its modulator STOML3 (Poole et al., 2014b; Ranade et al., 2014; Wetzel et al., 2017). Indeed small molecule inhibition of STOML3 in the mouse forepaw reversibly reduces the ability of the mouse to perceive mechanical stimuli (Wetzel et al., 2017). It is known that the expression of mechanoreceptor specific potassium channels like KCNQ4 modulate the mechanoreceptor response to low frequency sinusoidal stimuli (Heidenreich et al., 2012), so it is conceivable that mouse forepaw afferents express a different complement of potassium channels than mechanoreceptors innervating other skin regions. The high sensitivity of mouse forepaw Meissner’s mechanoreceptors suggests that enhanced tactile acuity in the forepaw has been selected for during evolution and may have fitness advantages by allowing mice to more accurately select their dietary intake according its tactile properties. The physiological and anatomical specialization of forepaw Meissner’s corpuscle receptors was all the more remarkable as other mechanoreceptors, like SAMs and Aδ-nociceptors (Fig. 5), showed mechanosensitive properties that were invariant across the skin areas examined. Thus we can be confident that data gathered on such receptors can compared regardless of the skin area examined. Conversely, it is also clear from our data that RAMs from the forepaw cannot be compared with those recorded from the hind paw.

D-hair receptors are the most sensitive cutaneous mechanoreceptors and typically have large receptive fields and can be activated by almost all hairs within the receptive field (Brown and Iggo, 1967; Burgess et al., 1968; Lewin and McMahon, 1991; Ritter et al., 1991). These mechanoreceptors were first described in cat hairy skin and have been described extensively in rodents, but have essentially identical properties in primates and are also found in humans (Adriaensen et al., 1983; Perl, 1968). The development of D-hair receptors is controlled by multiple neurotrophic factors. The number of D-hair receptors that develop in the post-natal skin is controlled by Nerve growth factor (NGF) (Lewin et al., 1992; Ritter et al., 1991) and in the adult animal D-hair receptors require Neurotrophin-4 (NT-4) for survival (Stucky et al., 1998, 2002). It has long been known that D-hair receptors express high levels of TrkB the main receptor for Brain derived neurotrophic factor (BDNF) and NT-4 (Li et al., 2011; Rutlin et al., 2014; Shin et al., 2003). A novel physiological and anatomical feature of D-hair receptors is that their end-organ consists of an array of lanceolate endings that form a horseshoe around the innervated hair (Bernal Sierra et al., 2017; Rutlin et al., 2014) and this anatomical arrangement is thought to underpin strong direction sensitivity. Polarized expression of BDNF in the developing follicle is necessary for the asymmetric structure of the lanceolate endings and for direction sensitivity of the receptor. Here we describe a unique population of small hairs in the mouse hind paw glabrous skin that appear to be predominantly innervated by D-hair receptors (Fig. 6). We could confirm that these D-hair receptors are also directionally sensitive being best activated as the hair was moved against the skin (Fig. 6). In the hairy skin of the mouse back D-hair receptors were described as best activated by movement of the hair in the rostral direction which corresponded with the hair shaft pulling away from the lanceolate endings. Caudal movement of the hair would result in the shaft pushing against the main lanceolate array (Rutlin et al., 2014). For the D-hair receptors recorded in the glabrous skin the situation was directly the opposite, thus the receptors were most sensitive in the direction where the shaft would be pushed against the main lanceolate array (Fig. 6). There is evidence that protein tethers may link transduction channels in sensory neurons to the extracellular matrix (Chiang et al., 2011; Hu et al., 2010) and such tethers could link lanceolate endings to the hair follicle shaft (Li and Ginty, 2014). It is conceivable that tethers that link the hair shaft and transduction channels in the lanceolate endings are pulled to gate channels in D-hair receptors in the mouse back skin. The opposite directional sensitivity with the same anatomical arrangement in glabrous skin D-hair receptors means that this model should be modified. It is, for example, conceivable that a tether based transduction mechanism is used by both receptors, but it is the precise positioning of the tethers that determines direction sensitivity. It will only be possible to resolve this issue definitively when the molecular identity of the tether and its relationship to the transduction channels is known. Nevertheless, the unique physiological properties of the glabrous skin D-hair receptor suggest that these hairs serve a distinct and specific sensory role in comparison to the hairy skin D-hair receptors. Recently, several reports in mice have suggested that it is D-hair receptors that drive touch evoked pain under neuropathic conditions (Dhandapani et al., 2018; Peng et al., 2017; Ventéo et al., 2016). Since in neuropathic models behavioral assessments of increased sensitivity to mechanical stimuli are made using von Frey hairs applied to the glabrous skin of the hind paw, it is likely that glabrous skin D-hair receptors drive neuropathic behavior. The role of hairy skin D-hair receptors and the glabrous skin D-hair receptors in normal touch behavior remains unresolved. Experiments in humans have shown that activity in some mechanoreceptor units e.g. RAMs and SAM type I receptors are capable of triggering conscious perception of touch (Ochoa and Torebjork, 1983; Sanchez Panchuelo et al., 2016; Torebjork et al., 1987; Vallbo, 1981). However, activity in other mechanoreceptors like SAM type II receptors does not appear to trigger conscious touch perception (Ochoa and Torebjork, 1983). It is thus entirely possible that under normal circumstances activity in D-hair receptors provides sensory input that modulates motor behavior but does not contribute to tactile perception. We show here that glabrous D-hair receptors are not ubiquitous in rodents, but are probably evolutionarily ancient. The appearance of these receptors in rodent species that diverged more than 65 million years ago (Fabre et al., 2012) suggests that there may be a developmental program leading to their appearance that has been switched on or off during speciation. Rodent species that possess glabrous D-hair receptors like the Grasshopper mouse or the Damaraland mole-rat occupy totally different habitats and it is not obvious what might have provided selective pressure for the retention of glabrous D-hair receptors.

In summary, we have provided evidence for unique mechanoreceptor specializations in the mouse glabrous skin. Our data provide a solid basis to evaluate how rodents use their glabrous skin surfaces to explore their tactile environment.

## Acknowledgments

This work was supported by grants from the Deutsche Forschungsgemeinschaft (SFB 665 Project B6 to GRL) and a European Research Council advanced grant (ERC 294678). We thank Alison Barker for her comments on the manuscript. We also acknowledge the technical assistance of the Advanced Light Microscopy technology platform of the MDC.

## Materials and Methods

### Ex vivo skin-nerve preparations

Electrophysiological recordings from fibers of the saphenous nerve were done using an *ex vivo* skin nerve preparation as previously described (Koltzenburg et al., 1997; Milenkovic et al., 2008) with some minor modifications. The animal was sacrificed by cervical dislocation and the hairs of the limb were shaved off. The hairy skin of the upper leg was carefully removed and the saphenous nerve was dissected up to the hip and cut. The paw skin with the connected nerve was transferred into a bath chamber and fixed with insect needles using the outside out configuration (Fig. 1C). The bath chamber was constantly perfused with warm (32°C) oxygen-saturated synthetic interstitial fluid (SIF). The nerve end was passed through a narrow channel into an adjacent recording chamber that was filled with mineral oil. Since the outside out configuration was used, a 1ml pipette was used to regularly flush the dermis of the skin sample with fresh SIF buffer.

For the tibial nerve *ex vivo* skin nerve preparation the hairy skin of the hind paw was removed, the bones detached, and any remaining muscle and ligament tissue was carefully dissected (Milenkovic et al., 2014). We noticed that remaining muscles or ligaments around the nerve trunks decreased the duration with which the preparation could be used and often interfered with mechanical and electrical stimulation procedures. Therefore, as much muscle and tendon tissue as possible was removed. A clean preparation (Fig. 1D) allowed for several hours of recordings with reliable mechanical and electrical stimulation using the outside out configuration with bath conditions as described above.

For the forepaw *ex vivo* skin nerve preparation, we removed the hairy skin of the forepaw, the bones, ligaments and muscle tissue instead of stripping the glabrous skin from the forepaw. Due to the challenging anatomy around the thenar and hypothenar pads some remaining muscle tissue was not dissected. Using the outside out configuration (Fig. 1I) allowed for high quality recordings of several hours even though the total experimental times were shorter compared to experiments using the hind paw preparations described above (approximately 2-3 hours shorter).

Single unit recordings were made as previously described (Koltzenburg et al., 1997; Milenkovic et al., 2008, 2014). Fine forceps were used to remove the perineurium of the nerve. Fine nerve bundles were teased and attached against to a platinum wire that served as the recording electrode. Mechanical sensitive units were first located using blunt stimuli applied with a glass rod. The spike pattern and the sensitivity to stimulus velocity were used to classify the unit as previously described (Milenkovic et al., 2008; Ranade et al., 2014). A Powerlab 4/30 system and Labchart 7.1 software with the spikes-histogram extension were used to record the raw data. The conduction velocity was measured from the latency between the electrical stimulus and arrival of the action potential at the electrode. All mechanical responses analyzed were corrected for this time delay. The total distance that the action potentials traveled could be measured by taking the distance between the stimulation electrode (receptor site) and the recording electrode (nerve end). The conduction velocity (CV) could be measured using the formula CV = distance / time delay, with CVs > 10m/s were classified as Aβ, < 10m/s as Aδ and < 1.5m/s as C-fibers.

Mechanical stimulation of receptor units were performed using a piezo actuator (Physik Instrumente, P-841.60; see Fig. 2A) and a double-ended Nanomotor (Kleindiek Nanotechnik, MM-NM3108) connected to a force measurement device (Kleindiek Nanotechnik, PL-FMS-LS). A magnetic stand connected to a micromanipulator assisted in the positioning of the piezo actuator. As the different receptor units are tuned to specific stimuli, different mechanical stimuli were used based on the unit type: A vibrating stimulus with increasing amplitude (using the piezo actuator) was used with a vibration frequency of 20Hz (Fig. 2B). The force needed to evoke the first action potential was measured. Only the low threshold mechanoreceptors were tested with this stimulus. A dynamic mechanical stimulus with a ramp and hold waveform was used with a constant force (using the piezo actuator; average force of 100mN) and repeated with varying probe movement velocity (0.075mm/s, 0.15mm/s, 0.45mm/s, 1.5mm/s). Only the spikes of the dynamic phase were analyzed, and only low threshold mechanoreceptors were tested with this stimulus (see Fig. 2C). A static mechanical stimulus with a ramp and hold waveform was used with a constant fast ramp (1.5-2mN/ms) and repeated with varying amplitude (using the double-ended Nanomotor). Only spikes evoked during the static phase were analyzed (Fig. 2D). This stimulus was only used to investigate the slowly adapting SAs and AMs. Single hair stimulation: To selectively move single hairs a fine glass capillary (tip size approximately 150μm) attached to a Micromanipulator (Kleindiek Nanotechnik, MM3A) was used. Single hairs could be mounted inside the capillary (Fig. 5E) and then displaced into 2-Dimensional space using a step protocol (8, 16, 32, 64 and 128 course steps, frequency of 3200 steps/second, step-size approximately 6.3μm – this slightly varies depending on the angle, total length of the axis and distance to the skin). The action potentials triggered during the probe movement were quantified. Pictures and Videos were visualized and captured using a binocular with a camera attached (Leica, MS5, IC80, LAS software). The video signal of both screens displaying the camera signal and the electrophysiological responses were simultaneously recorded (DebugMode, Wink) and saved as single video frames.

### Immunofluorescence and anatomy

Skin tissue was dissected removing the hypodermis, ligaments and muscle tissue and stretched out using insect pins. The samples were then fixed at room temperature (RT) for 45-60min. in PFA (4%). For whole mount staining the samples were bleached at 4°C for 24h in 10% H_2_O_2_, 18% DMSO and 72% methanol, washed five times with methanol and post fixed at 4°C for 24H in 20% DMSO, 80% methanol.

Four to six week old heterozygous Cav3.2Cre mice were anesthetized by an intraperitoneal injection of ketamine (100mg/kg) and xylazine (10mg/kg). The sciatic or saphenous nerve were exposed and 2 μl of AAV-flex-tdTomato (AV2/9.CAG.FLEX.tdTomato.WPRE.bGH; Penn Vector Core, PA, USA) with a titer of 3.71 E+12 was slowly injected into using a pulled glass capillary attached to a Hamilton microliter syringe. After the injection, the capillary was left in place for additional three min.

For gelatin vibratome sections the tissues were placed into disposable embedding molds (Polysciences, T-8) filled with warm (45°C) 20% gelatin dissolved in PBS and positioned for sagittal sectioning until curing. The gelatin block was post fixed at 4°C ON in 4% PFA and cut in PBS into 120μm thick slices using a vibratome (Leica, VT100S).

### Tissue clearing

The skin samples were washed in PBS and incubated at 4°C for 1h with blocking solution (PBS + 5% goat or donkey serum and 0.4% Triton X-100 (Sigma)). Primary Antibodies were incubated for 48h at 4°C in blocking solution. After thorough washing with PBS the secondary antibodies diluted in blocking solution were incubated for 24h. Samples were washed in PBS and incubated each time at 4°C for 12h in a mix of 1:1 PBS/H2O and a rising concentration of 2,2’-Thiodiethanol (TDE, Sigma Aldrich). TDE concentrations were increased from 10%, 25%, 50% and 97% at which the samples were stored and mounted onto slides and coverslips (remaining in a 97% TDE solution).

### Antibody staining

Antibodies used were rabbit anti-S100 (Dako, 1:1000), chicken anti-NF200 (Millipore, 1:1000), Anti-rabbit Alexa Fluor 488 and anti-chicken Alexa Fluor 647 (both Invitrogen, 1:800).

### Image acquisition and analysis

Tiled image stacks were taken using a confocal microscopes (LSM 700 and LSM710 Carl-Zeiss) running Zen 2009 software was used. Nerve fibers innervation of Meissner’s corpuscles were visualized and counted using Fiji/ImageJ with Bio-Formates extension. Only nerve fibers labeled for both S100 and NF200 were counted. The volume of single Meissner’s corpuscle and the volume between the basement membrane and the dermis (stratum basale, stratum spinosum, stratum granulosum) of each stack were estimated using the integrated area tool of Fiji/ImageJ multiplied by the stack thickness (usually 2μm). Total volume was calculated by the sum of the corresponding stack volumes.

### Electron Microscopy

Saphenous, tibial, median and ulnar nerves were dissected and fixed in 4% paraformaldehyde and 2.5% glutaraldehyde in phosphate buffer, postfixed and contrasted with osmium tetroxide (Garratt et al., 2000) and embedded in Technovit 7100 resin (Heraeus Kulzer, Wehrheim, Germany). Myelinated nerve fibers were quantified using thin toluidine blue stained 1μm semi-thin sections (Figure 4G).

### Additional Software used

Carl-Zeiss Zen 2009 v2.3 lite, Fiji/ImageJ and Adobe creative suite v5.5 were used for Image, video and figure arrangement and processing. Raw data was stored and processed using Microsoft excel. Statistical tests were performed using GraphPad prism5 and 6.

## Supplementary Files Supplementary Figure 1

Rodent species that lack glabrous D-hair receptors: **A** One North American rodent species that lacks glabrous D-hair receptors. **B-K** Ten African rodent species that clearly ack glabrous D-hair receptors.

**Supplementary Video 1**

A maximum intensity projection of one entire foot pad showing NF200-positive fibers innervating Meissner’s corpuscles.

**Supplementary Video 2**

The video shows the experimental set-up for recording directional sensitivity of glabrous D-hair receptors. The single hairs is moved within a glass and single-unit firing behavior is shown in response to 4 axis of movement.

**Supplementary Video 3**

Maximum intensity projection of a tdTomato labeled endings around a single glabrous hair follicle.

